# SynDRep: A Knowledge Graph-Enhanced Tool based on Synergistic Partner Prediction for Drug Repurposing

**DOI:** 10.1101/2024.08.13.607713

**Authors:** Karim S. Shalaby, Sathvik Guru Rao, Bruce Schultz, Martin Hofmann-Apitius, Alpha Tom Kodamullil, Vinay Srinivas Bharadhwaj

## Abstract

**Motivation:** Drug repurposing is gaining interest due to its high cost-effectiveness, low risks, and improved patient outcomes. However, most drug repurposing methods depend on drug-disease-target semantic connections of a single drug rather than insights from drug combination data. In this study, we propose SynDRep, a novel drug repurposing tool based on enriching knowledge graphs (KG) with drug combination effects. It predicts the synergistic drug partner with a commonly prescribed drug for the target disease, leveraging graph embedding and machine learning techniques. This partner drug is then repurposed as a single agent for this disease by exploring pathways between them in KG.

**Results:** HolE was the best-performing embedding model (with 84.58% of true predictions for all relations), and random forest emerged as the best ML model with an ROC-AUC value of 0.796. Some of our selected candidates, such as miconazole and albendazole for Alzheimer’s disease, have been validated through literature, while others lack either a clear pathway or literature evidence for their use for the disease of interest. Therefore, complementing SynDRep with more specialized KG, and additional training data, would enhance its efficacy and offer cost-effective and timely solutions for patients.

**Availability and Implementation:** SynDRep is available as an open-source Python package at https://github.com/SynDRep/SynDRep under the Apache 2.0 License.

## Introduction

Despite tremendous technological, regulatory, and scientific advances that increase the efficiency of drug research and development, the resulting therapeutic outcomes need to catch up with the corresponding spending on these advances (Ashburn and Thor 2004; Scannell *et al*. 2012). Additionally, the rising cost and time required to develop new drugs have resulted in lower profits for the pharmaceutical sector and a longer response time to disease outbreaks (Pushpakom *et al*. 2018). Conversely, drug repurposing, i.e., finding novel indications for current drugs, has advantages over de-novo drug development, including shorter development time and lower cost risk (Choudhury, Arul Murugan and Priyakumar 2022; Hua *et al*. 2022), since compounds already investigated and approved by regulatory bodies, incorporating safety and efficacy profiles, can be reassessed critically in a new therapeutic context (Lage-Rupprecht *et al*. 2022).

In recent years, drug repurposing research has greatly benefited from the exploding growth of biomedical databases. Therefore, plenty of computational techniques have been devised to analyze different biomedical data systematically to hypothesize new indications for a drug or to find new drugs for a specific disease (Jarada, Rokne and Alhajj 2020; Luo *et al*. 2021; Pan *et al*. 2022). Computational drug repurposing approaches are mostly data-driven; they encompass the systematic analysis of data from various modalities, e.g., chemical structure, proteomic data, gene expression, genotype, or electronic health records, which can then drive the repurposing hypotheses (Hurle *et al*. 2013; Zong *et al*. 2022). For practical analysis of such vast data types, measures for appropriately aggregating them in an informative manner need to be taken. One of these measures is the organization and representation of data into a knowledge graph (KG), which aids in identifying semantic connections between multiple resources and allows for knowledge reasoning (Chen, Cheng and Li 2020; Gao, Ding and Xu 2022). Extending these mechanistic KGs with drug-related data to form drug-target-mechanism-oriented data models results in so-called PHARMACOMES (Lage-Rupprecht *et al*. 2022).

Pharmacomes with their integration of pathophysiology mechanisms, drug targets, and drugs/compounds offer the possibility to look at dual targeting strategies and combinatorial targeting of different pathophysiology mechanisms through combinations of drug repurposing candidates. Drug combinations offer excellent efficacy in treating multifactorial diseases involving more than one genetic pathway, such as cancer (Zhou, Edil and Li 2023), diabetes (Dubourg *et al*. 2022), Alzheimer’s disease (AD) (Knorz and Quante 2022), and cardiovascular diseases (Lombardi *et al*. 2020). In principle, they also offer the option to specifically target co-morbidity pathways. Therefore, incorporating new links among drugs into pharmacomes, indicating drug combinations, paves the way for developing new synergistic drug combinations. It warns of potential drug-drug interactions in a more comprehensive way that depends on direct as well as indirect links between drugs. Applying some link prediction algorithms afterward will predict new drug relationships, gain more insights into drug mechanisms, and eventually repurpose drug candidates for various diseases.

For a long time, drug synergy studies depended on trial and error, which suffers from high labor and time costs and exposes patients to ineffective treatment or undesirable side effects (Pang *et al*. 2014; Day and Siu 2016). This was then replaced by high-throughput screening (HTS), where many measurements can be produced reasonably fast and at a lower cost (He *et al*. 2018). During HTS, different concentrations of two or more drugs are applied to a cell line. However, the high genomic correlation between the original tissues and the derived cell lines remains imperfect (Ferreira *et al*. 2013). Moreover, HTS cannot cover the whole combination space for drugs (Goswami *et al*. 2015). Computational methods such as deep and machine learning (DL/ML) models can efficiently explore the vast synergistic space using the available HTS synergy data. Recent methods range from systems biology (Feala *et al*. 2010), kinetic models (Sun *et al*. 2016), mixed integer linear programming (Pang *et al*. 2014), computational methods based on Drug-induced gene expression profile and dose-response curves (Goswami *et al*. 2015), to ML approaches including Random Forests and Naive Bayes methods (Li *et al*. 2015; Wildenhain *et al*. 2015), and DL approaches such as deep neural networks, graph autoencoder, and convolutional neural network (Preuer *et al*. 2018; Kuenzi *et al*. 2020; Sun *et al*. 2020; Kim *et al*. 2021; Liu and Xie 2021; Li *et al*. 2023). However, these methods are restricted to predicting synergistic combinations and not consider drug synergy prediction as an intermediate step in the drug repurposing process. In our approach, we leverage the synergistic prediction as a foundation for the repurposing process.

We propose a new drug repurposing tool (SynDRep), which depends on enriching knowledge graphs with drug combination effects. Our approach selects repurposing candidates, by predicting synergistic drug partners of a commonly prescribed drug for the target disease. This is followed by the selection of “safe drug partners” as a single-agent therapy for the disease. The drug’s candidacy for repurposing is confirmed by exploring the pathway within the KG between the drug and the target disease. Additionally, experimental evidence about the beneficial effect of the candidate on target disease supports the repurposing profile. Therefore, this approach combines the speed and cost reduction of the computational approach with the accuracy and certainty of manual curation and expands the current drug repurposing landscape with a new concept relying not only on drug-disease-target semantic connections but also on the drug-drug synergy effect.

## Methods

### 1. Data collection

The primary objective entailed the integration of drug-drug relationships into an established KG to serve as a foundational framework for subsequent computational methodologies (Figure 1).

**Figure 1.**
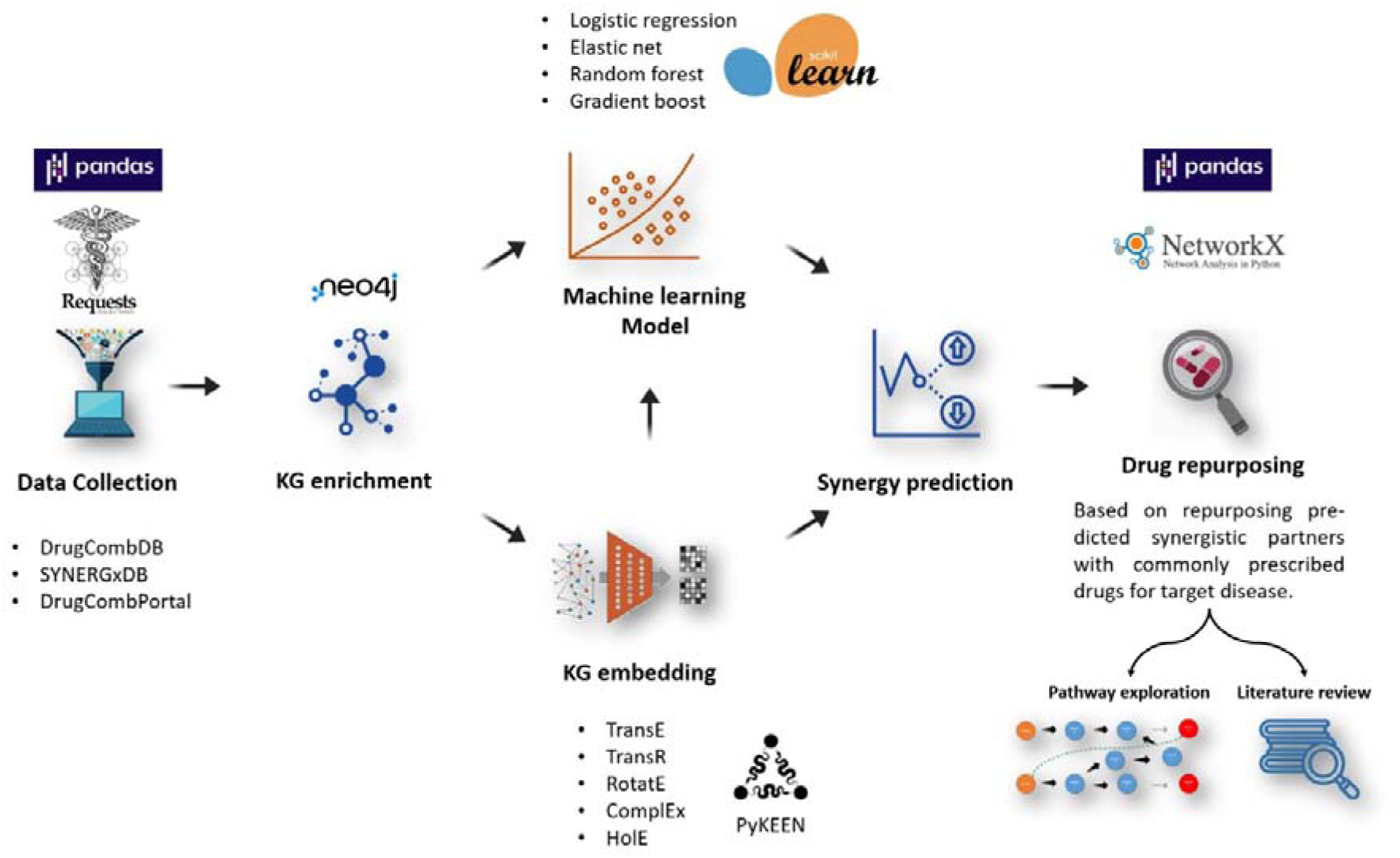
The overall workflow of the study, including the Python packages and data sources used. The work starts with data collection and refinement from combination databases using Pandas and Requests, then the synergy data is fed into a neo4j instance of the KG. Third, ML using scikit-learn, embedding using PyKEEN, or embedding followed by ML was used to model and predict the synergies. Finally, the identification of repurposing candidates by predicting synergistic drug partners for commonly prescribed drugs for the target disease and repurposing safe partners as a single agent for this disease. Candidate profiles are confirmed by examining the existence of pathways within the knowledge graph between the candidate drug and the target disease using Pandas and NetworkX. Experimental evidence from the literature supporting the candidate’s beneficial effect on the target disease further validates the repurposing profile.

Consequently, the initial phase involved the careful selection of a comprehensive KG such as the Human Brain Pharmacome (HBP) and appropriate sources for drug-drug combinations including drug synergy databases e.g., DrugcombDB, DrugcombPortal, and SYNERGxDB.

#### a. Human Brain Pharmacome

The KG selected for this study was the Human Brain pharmacome (https://graphstore.scai.fraunhofer.de/, by selecting the pharmacome option under Database) (Lage-Rupprecht *et al*. 2022), which is a comprehensive KG consisting of 136838 nodes and 731974 edges that combines knowledge from various sources with a focused drug-target-mechanism-oriented data model. It contains information curated from Bibliographic databases such as PubMed, Pathway databases such as Reactome, KEGG, and Pathway Commons, Protein-protein interaction databases such as IntAct, BioGRID, and StringDB, and Drug databases such as DrugBank, Clinical Trials, Sider, and ChEMBL. The data of this pharmacome has been extracted from the online source, stored locally using neo4j, and formed the base for the next step of KG enrichment.

#### b. Drug Synergy databases

There are many databases for drug combinations and their synergism or antagonism. To accommodate our expansive pharmacome we selected databases that contain the highest number of drugs. Drug combination effects have been gathered from DrugcombDB, DrugcombPortal, and SYNERGxDB (Zagidullin *et al*. 2019; Liu *et al*. 2020; Seo *et al*. 2020). The scores for synergism models, such as the highest single agent (HSA) model (Berenbaum 1989), Bliss model (Bliss 1939), Loewe model (Loewe 1953), and the Zero interaction potency (ZIP) model (Yadav *et al*. 2015), were used to supplement the new edges created in the next step. These models consider in their calculation the different effects of drug combinations at different drug concentrations.

### 2. KG enrichment

The KG enrichment was done over several steps. Data extracted from drug combination databases were deduplicated. The synergism scores from the same combinations with different scores were averaged. Some combinations of drugs produce synergism in one cell line and antagonism in another. Therefore, to remove the effect of different cell types, we selected only combinations that produced either synergism or antagonism on all cell types. Moreover, to go with a standardized approach, we selected only one synergism score (ZIP score), as the ZIP model encompasses the Loewe additivity and the Bliss independence. In addition, it is more accurate at detecting potency changes in drug combinations compared to HSA and Bliss independence models (Yadav *et al*. 2015).

### 3. Classical Machine learning

In order to predict new synergistic relations between drugs, we started with the classical ML approaches, to assess their ability and efficiency for link prediction compared to KG embedding. Four ML models, namely: logistic regression, elastic net, gradient boosting, and random forest, were selected along with the features of each pair of drugs to classify their combination either into synergism or antagonism. These features encompassed aspects related to the KG, as well as physicochemical attributes of the drugs as depicted in Table 1. KG features, which depend on the network structure and topological features were extracted from the pharmacome using NetworkX (Hagberg, Schult and Swart 2008), a Python package for the creation and study of the structure of complex networks. Physicochemical attributes of the drugs were extracted from PubChem or computed using the RDKit Python package (Landrum 2023). The features, labels, and models used are listed in Table 1. The classification was carried out over 10-fold cross-validation using Grid Search optimizer, and the model performance was assessed based on areas under the receiver operating characteristic curve (ROC-AUC) values for all the models. Data was first split into training (80%) and hold-out test set (20%). The cross-validation loops were performed by further splitting of the training data into 80% training and 20% validation for hyperparameter optimization (HPO). Following HPO, 10 model instances were trained using the best parameters and evaluated on the held-out test set using the ROC-AUC score.

**Table 1.**
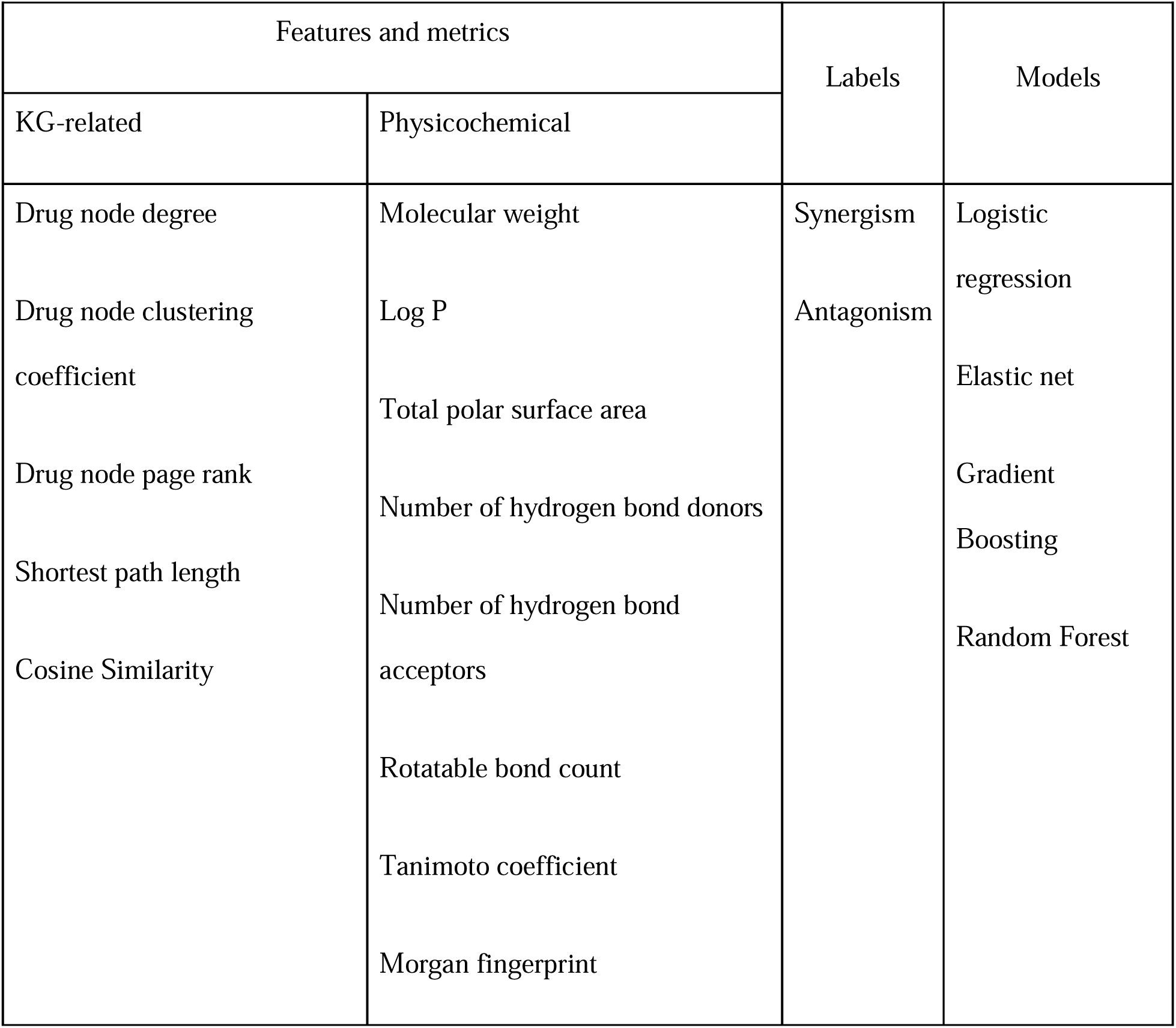
The features and labels used to train the different ML models.

Following a comprehensive assessment of all models and the calculation of ROC-AUC, we utilized the elastic net model to predict the synergistic interactions between each pair of drugs in the pharmacome. Subsequently, combinations predicted as synergistic were chosen to constitute the predicted synergism set. A literature check was conducted to validate the top five synergistic combinations based on their predicted probabilities during the initial prediction process. This validation step aimed to ensure the credibility and accuracy of the model’s predictions by cross-referencing them with existing scientific literature.

### 4. KG Embedding

To embed the enriched pharmacome, we used PyKEEN (Python KnowlEdge EmbeddiNgs) (Ali *et al*. 2021), a Python package designed for training and evaluation of KG embedding models. We worked under stochastic local closed world assumption (SLCWA), where a randomized subset is drawn from the combination of head and tail generation strategies, initially defined in local closed world assumption, and these selected triples are treated as negatives. This approach offers several advantages, including the lower load of computation and the flexibility to include new negative sampling strategies.

#### a. Data splitting

To prevent overfitting, the set of triples obtained from the extraction of enriched pharmacome was then stratified using the PyKEEN into a training set (80%) and a test set (20%). To prevent the dissemination of the test set into the training set during the HPO or training of the model, we isolated the test set, and the training set was further split into training (80%) and validation (20%) sets. We checked that each split contained the corresponding percentage of triples and that the training set contained all the relation types in the pharmacome to ensure that the test and validation sets did not contain any relation type new to the model after training. To further assess the model’s efficiency in predicting drug-drug relations, one more test set was formed from the original test set, the drug-drug test set, which contained only the drug-drug relations from the test set. These two sets were used to evaluate model performance.

#### b. Model selection

We selected five models for embedding the pharmacome: TransE, TransR, RotatE, ComplEx, and HolE. We afterward used the best-performing model to predict the new synergistic or antagonistic relations. Predictions were made using the two entities as head and tail, and the model predicts the relation type between them. The output of the prediction model is a ranking of the possible relations between these two entities according to a score produced by the model. Therefore, we selected the first three predictions to assess the model’s performance by calculating the percentage of True prediction in each rank compared to all predictions in this rank (equation):

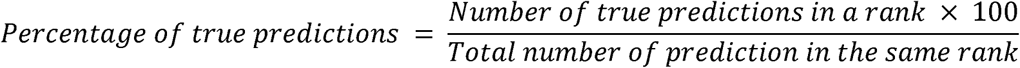

We calculated the multi-class ROC-AUC using the highest-ranked prediction for each pair of drugs in the test set. This involved converting the actual and predicted relation types into binary form and then averaging the ROC-AUC values for all relation types, as shown in the following equation:

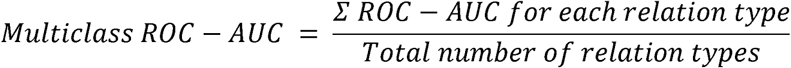

Based on the values of the percentage of true predictions and multi-class ROC-AUC, we selected the best-performing model for further prediction of the drug-drug relations that are not in the pharmacome.

### 5. Synergy prediction

After assessing all models and calculating the percentage of true prediction, we selected RotatE to predict all drug-drug relations further. Utilizing the trained RotatE model, we predicted the relations between each drug pair within the pharmacome. Both the forward case (with drug A as a head and drug B as a tail) and the reverse case (with drug B as a head and drug A as a tail) were predicted. The predicted relationship which ranked as the first was then extracted for each drug pair. To refine our dataset, we eliminated cases where nonmutual synergism or antagonism was observed. Then, we segregated the predicted dataset into synergism and antagonism categories, focusing on selecting data that showed synergistic interactions for subsequent in-depth analysis. A comprehensive literature review was conducted to validate the model’s predictive power, particularly for the predictions with the highest scores.

### 6. Drug repurposing

The synergy effect frequently stems from different drugs having influences on the same, parallel or even different pathways essential for an observed phenotype, and synergy is induced by targets aggregating at specific pathways that control the state of the disease (Cokol *et al*. 2011; Chen *et al*. 2015). Consequently, if the predicted synergy combination has a shared pathway in our pharmacome, then it is highly probable that the synergistic partner to a commonly prescribed drug for the target disease can be used solely for the management of this disease. Therefore, we assessed the predicted synergistic combination that includes drugs prescribed for AD, schizophrenia, and bipolar disorder to determine the plausibility of repurposing their synergistic partner for these diseases. Based on our predictions, we selected a list of drugs that exhibited the highest-scoring synergistic combinations for each drug. It is noteworthy that our selection criteria excluded drugs with cytotoxic or severe side effects, such as anti-cancer or carcinogenic drugs, ensuring that the chosen repurposing candidates prioritize safety considerations. This approach was initially followed by a meticulous search for a possible common pathway in the pharmacome between the two drugs in the combination and the disease. Subsequently, we reviewed the literature for possible studies about using these repurposed candidates as single agents for the disease of interest.

### 7. Causal-only pharmacome

Due to the lack of clarity in certain relations within the pathways explored in the HBP between repurposing candidates and the target diseases, we repeated the entire trial using a causal-only version of the HBP. In this version, we retained only the relations indicating direct causality between entities, such as increases, decreases, or causes no effect. Additionally, the presence of hubs within the pharmacome’s structure, primarily stemming from disease nodes, could introduce their overrepresentation during embedding and ML analysis. Consequently, these nodes and their connecting relationships were also removed before KG embedding. The disease nodes were removed before embedding, but we retained them in another copy of the causal-only pharmacome used for pathway confirmations. However, the performance of KG embedding models, in predicting drug-drug relations, was found to be suboptimal. Therefore, we introduced an additional step. We extracted vector embeddings of the KG and utilized them as inputs for training and testing ML models. By leveraging the best-performing ML model, we obtained the final predictions of drug-drug relations, which were then used for drug repurposing. We further challenged our repurposing approach for COVID-19, which has no node in our pharmacome, to explore the capabilities of our model, as a fast tool for drug repurposing for disease outbreaks.

## Results

### 1. KG enrichment

The KG enrichment was done over several steps as described in Methods section. The drug combination dataset contained 23171 pairs from 882 unique drugs, which were used to enrich the pharmacome. These combinations were further converted into edges and then added to the neo4j instance of the pharmacome by converting the values of ZIP scores into synergistic, antagonistic, or additive effects. To avoid class imbalance and model overfitting due to the extremely low number of additive effect relations, additive edges were not been added to the pharmacome. The enriched pharmacome was then completely extracted as triples of source, relation, and target and was used to train and validate ML and KG embedding models.

### 2. Classical machine learning

The cross-validation and synergy prediction results from the four selected ML models are detailed in Supplementary Results section 1. Due to the lack of virtually validating studies and because this classical ML approach does not take relation type into consideration, we conducted a graph-embedding-based approach, as explained in the next section.

### 3. KG Embedding

Lacking a virtual validation by confirming literature, we turned our focus to novel combination prediction using graph embeddings. To do so we performed pharmacome embedding using different algorithms to model and predict novel drug-drug links in the KG. When tested on the test set, RotatE model consistently outperformed other models in producing true predictions at the lowest rank (Figure 2). Based on these results, the RotatE model was selected as the model of choice for predicting relationships between drugs that do not have a direct link within the pharmacome.

**Figure 2.**
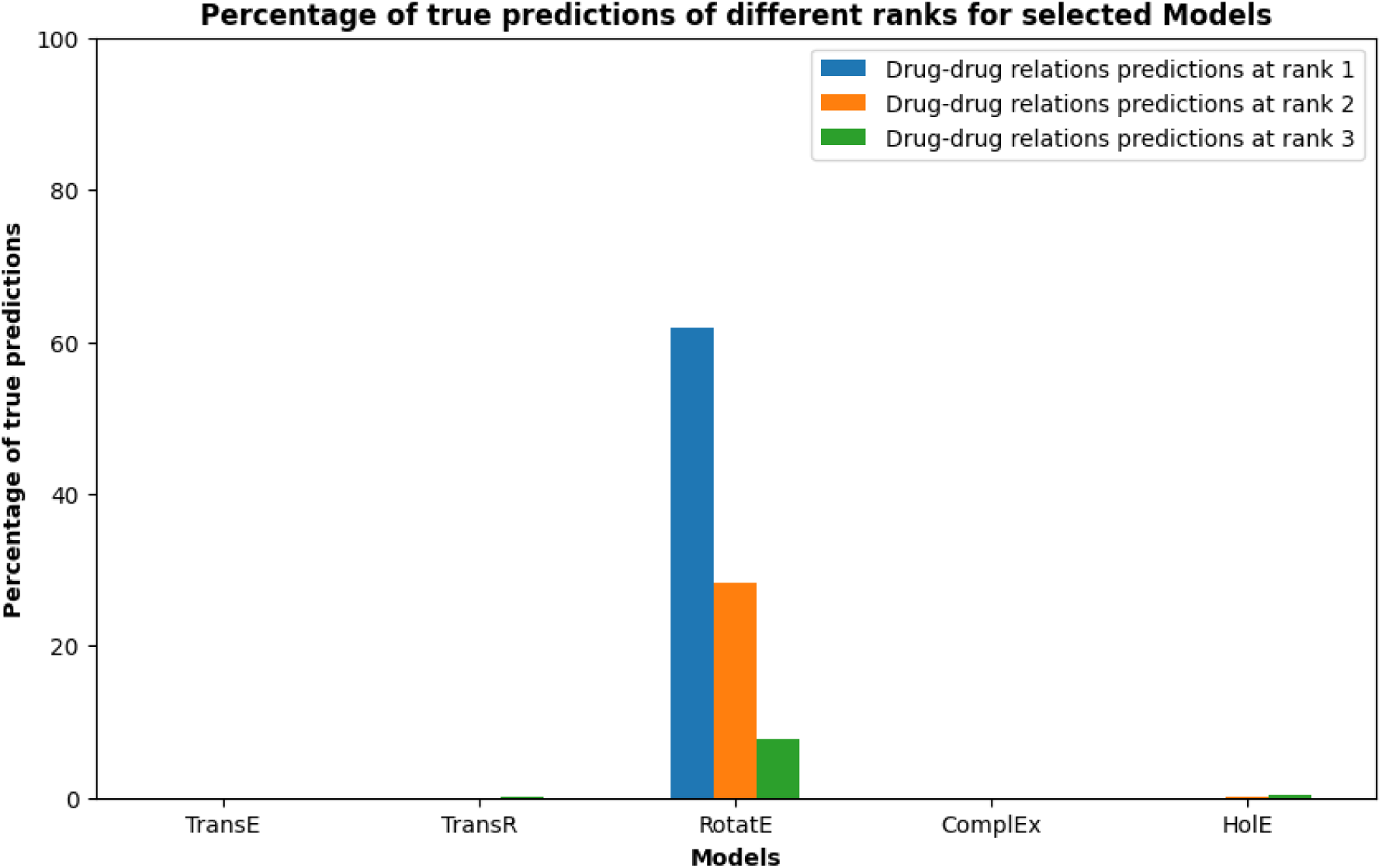
Percentage of true drug-drug relation predictions at different ranks for selected models. Optimum models were used to predict the drug-drug relations in the test set from the original HBP. Then, the predicted relations were compared to the actual relations to calculate the percentage of true predictions.

### 4. Synergy prediction

Synergy predictions were generated leveraging the trained RotatE model. The prediction set was further processed as outlined in Supplementary Results section 3. The five highest-scoring synergy combinations were subjected to a thorough literature review to validate the reliability of the model’s predictions, as detailed in Supplementary Table S2. The findings revealed that most of these combinations either exhibited documented synergy or were being utilized in combination for the treatment of the specific diseases for which they were intended.

### 5. Drug repurposing

The assessment of the plausibility of repurposing candidates for the target disease was done by revising the literature for scientific data about their use in the selected disease and determining the common pathways between the synergistic drugs and the disease in our pharmacome using the Python package NetworkX. This comprehensive approach enhances our understanding of potential therapeutic applications and facilitates informed decision-making regarding drug repurposing candidates. Detailed explanations of these candidates are elaborated in Supplementary Results section 4.

Leveraging a broad and highly connected KG such as HBP, with both casual and non-causal relations like association and complexity, can lead to suboptimal model training and prediction, as well as to less explainable pathways between drugs and diseases. In addition, the presence of nodes with high degrees in KG will lead to inadequate training and inaccurate predictions as well. We believe these factors contributed to the discrepancy between our model’s predictions for schizophrenia repurposing candidates and existing literature evidence. To address this, we conducted a subsequent trial using a causal-only pharmacome, where we removed non-causal relationships and hub nodes. The hub nodes were removed before embedding, but we retained them in another copy of the causal-only pharmacome used for pathway confirmations.

### 6. Causal-only pharmacome

After the removal of non-causal relations and disease nodes that form hubs in the HBP, the same KG embedding models were used to model the causal-only version of the HBP. HolE was the best model to produce true predictions at the lowest rank (74.90% for all relations), as shown in Supplementary Figure S1. Although it was also the best-performing model for drug-drug relations, the percentage of true predictions at the lowest rank was low (54.65%), indicating nearly random predictions (Supplementary Figure S2). Therefore, we changed the design of the experiment to incorporate KG embeddings, followed by ML model training and prediction using the extracted embedding vectors as input features. In this approach, we utilized the vector embeddings of the causal-only pharmacome without enrichment with drug-drug relations. The synergistic data was then used as labels for training and testing the ML models. In this run, HolE again proved to be the best-performing embedding model (84.58% for all relations), and random forest emerged as the best ML model with an ROC-AUC value of 0.796 (Figure 3 and Figure 4). We have made the causal-only pharmacome, enriched with drug-drug relations (predicted and from databases), available at: https://doi.org/10.5281/zenodo.12806409. Consistent with prior methods, we validated the top-scoring synergistic combinations through a literature review (detailed in Supplementary Table S3). This analysis found that most combinations (three out of five) either demonstrated synergy between drug pairs or one of the drugs enhanced the effect of the other.

**Figure 3.**
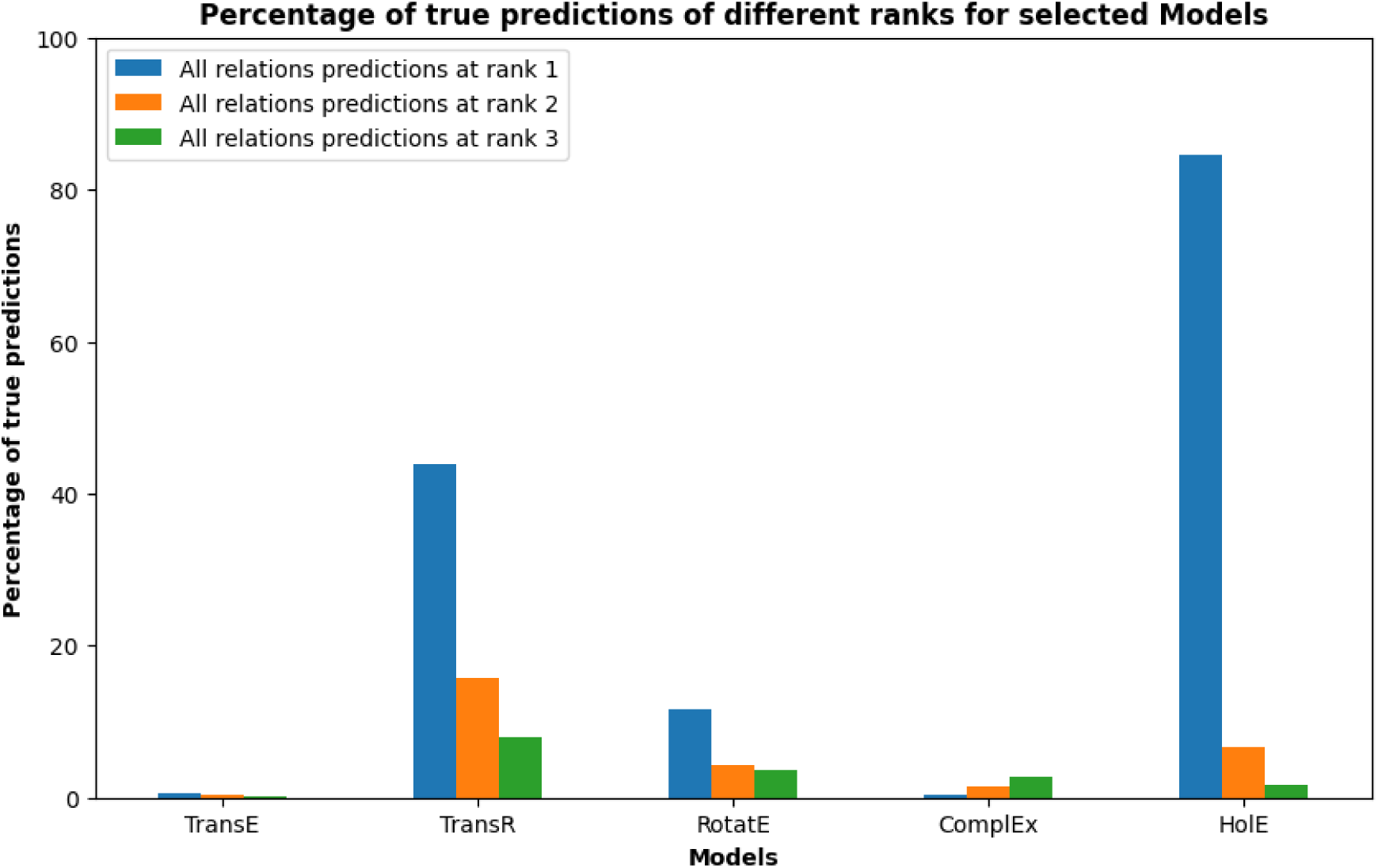
Percentage of true all relations predictions at different ranks for selected models used to embed Casual-only pharmacome before ML. Optimum models were used to predict the test set, and then the predicted relations were compared to the actual relations to calculate the percentage of true predictions. Embedding vectors of HolE were extracted and used as input features for the training and testing of ML models.

**Figure 4.**
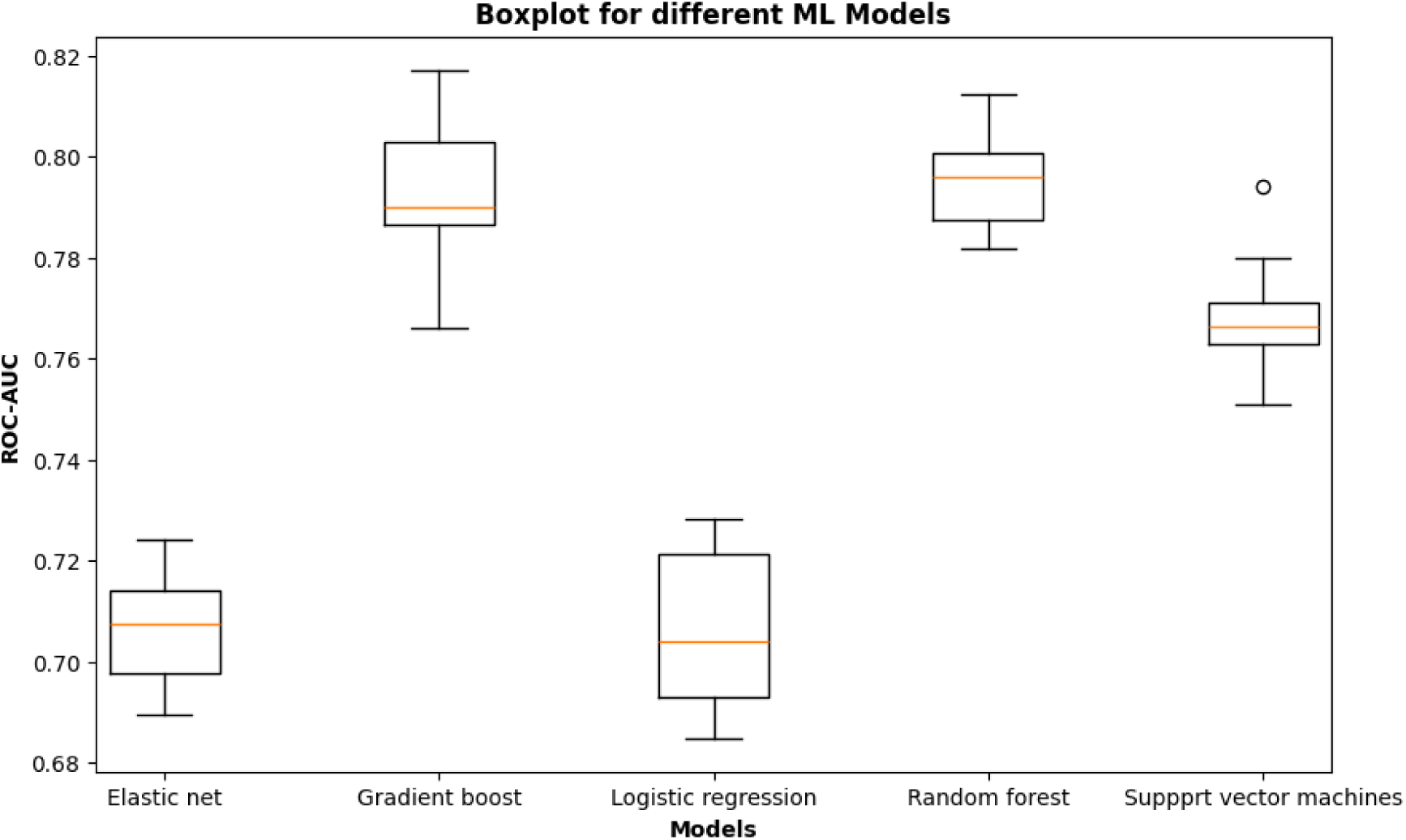
Benchmarking of ML models trained to classify between synergism and antagonism using the embedding vectors from causal-only pharmacome. Each boxplot shows the distribution of the ROC-AUC values over ten repeats of the ten-fold nested cross-validation procedure.

Subsequently, we selected safe drugs predicted to be synergistic partners with the previously chosen drugs for AD and schizophrenia, or bipolar disorder.

#### a. Alzheimer’s candidates

Based on our research, miconazole and albendazole have emerged as promising candidates for repurposing to treat AD. They were predicted to act synergistically with three and two of the selected AD drugs, respectively. By tracing their pathways to AD in the copy of the causal-only pharmacome, where we retained the disease nodes, we found that they share pathways with their synergistic partners to AD.

Miconazole is a broad-spectrum antifungal with some antibacterial activity (Wishart *et al*. 2018). On the other hand, it offers a potential therapeutic approach for early intervention in AD by promoting myelination of the medial prefrontal cortex and ameliorating neuroinflammation-mediated AD progression in different mice models (Yeo *et al*. 2020; Wang *et al*. 2022). A prominent common pathway of miconazole with donepezil, rivastigmine, and galantamine to AD was found in pharmacome, as shown in Figure 5. On the other hand, Albendazole is primarily employed as an anthelmintic to treat helminth infections (Sungkar *et al*. 2019). However, research on H4 neuroglioma cells has shown that albendazole can reduce Tau levels, suggesting a beneficial effect on AD (Dickey *et al*. 2006). In the causal-only pharmacome, it has a shared pathway with donepezil and rivastigmine, in which it intersects with them in decreasing the levels of phosphorylated Microtubule-associated protein tau (MAPT), which is a hallmark of AD (Figure 5). Comprehensive description of miconazole and albendazole pathways is elaborated in Supplementary Results section 5.

**Figure 5.**
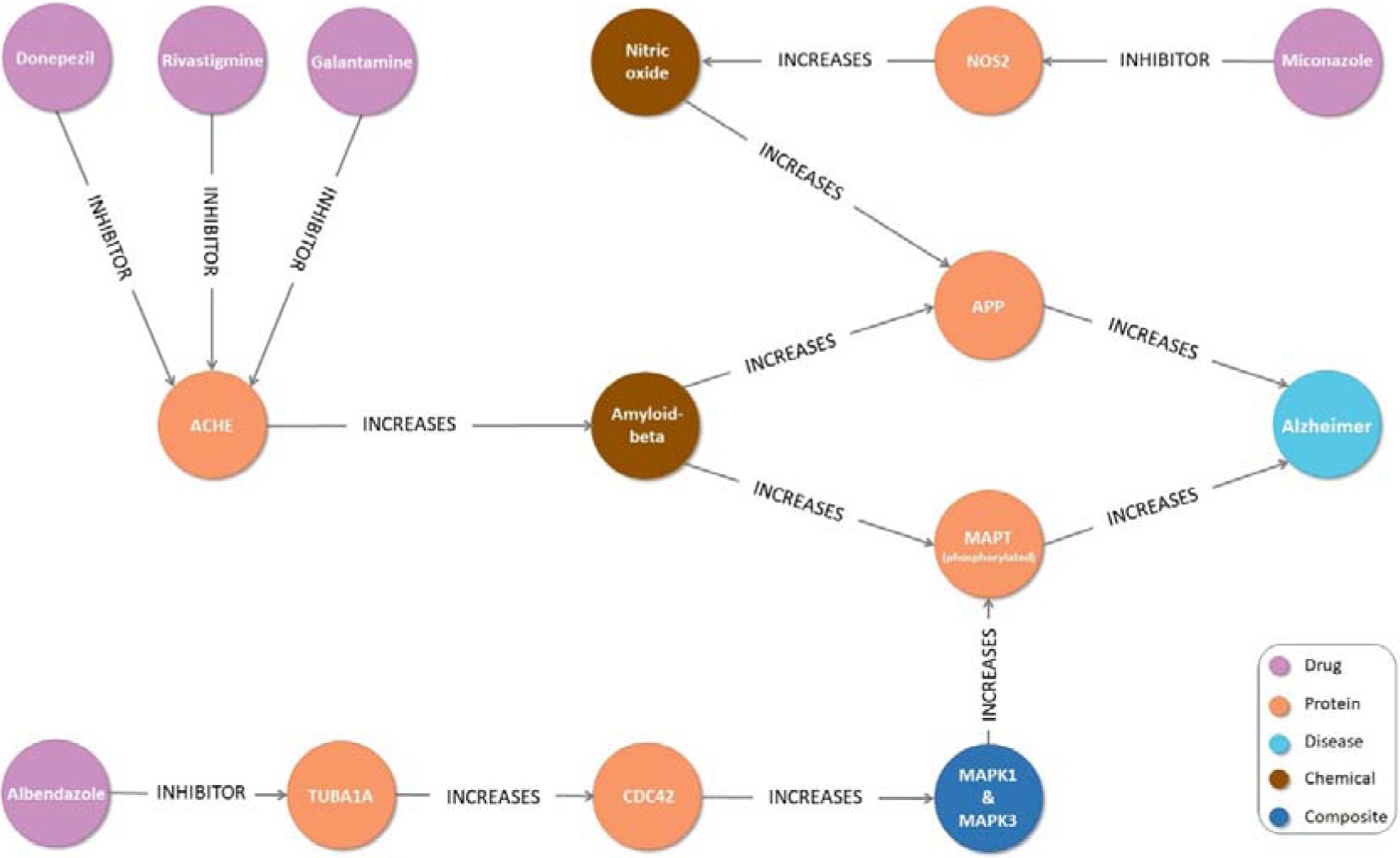
The shared pathways of miconazole, albendazole, donepezil, rivastigmine, and galantamine to Alzheimer’s disease. Disease nodes were retained in a copy of the causal-only pharmacome used for pathway confirmations. (ACHE: acetylcholinesterase, NOS2: nitric oxide synthase 2, APP: amyloid-beta precursor protein, MAPT: microtubule-associated protein tau, TUBA1A: tubulin alpha-1A protein, CDC42: cell division control protein 42 homolog, MAPK1 and MAPK3: mitogen-activated protein kinase 1 and 3).

Other candidates, such as disulfiram, auranofin, and finafloxacin, were also predicted as synergistic partners with AD drugs and studies, in cell and animal models, showed their beneficial effects for the management of AD (Madeira *et al*. 2012; Roder and Thomson 2015; Reinhardt *et al*. 2018; Upīte *et al*. 2020; Jun and Fang 2021; Guo *et al*. 2022). However, no supporting pathway in the pharmacome could be detected for these drugs. Additionally, prochlorperazine was consistently predicted as a synergistic partner with the three selected AD drugs. However, it has anticholinergic properties that in higher doses might worsen AD-associated dementia (Obara *et al*. 2019). Moreover, the explored pathways between prochlorperazine and AD were controversial, with some suggesting it might have a beneficial effect for the management of AD while others suggest it might exacerbate the condition.

#### b. Schizophrenia and bipolar disorder candidates

Upon detecting repurposing candidates for schizophrenia or bipolar disorder, we found no connecting pathways between any drug in the causal-only pharmacome and these diseases, even for the drugs that are typically prescribed for these conditions. Therefore, we couldn’t identify any repurposing candidates for schizophrenia or bipolar disorder based on the causal-only pharmacome.

#### c. COVID-19 candidates

We wanted to challenge our model further by checking its ability to predict synergistic drug combinations for diseases, which the KG was not built for, to test its ability to be used as a fast tool for repurposing drugs to new disease outbreaks. We selected a drug, baricitinib, used for COVID-19 and extracted the synergistic combinations from those predicted by our model. From these combinations, Prochlorperazine and disulfiram were identified as repurposing candidates based on their predicted synergy with baricitinib and their beneficial effects in COVID-19 management. These effects include the inhibition of SARS-CoV-2 entry by targeting the spike protein and ACE2, which were confirmed computationally by molecular docking and experimentally in VeroE6 and HEK293T-hACE2 cell cultures (Chen *et al*. 2022; Liang *et al*. 2023).

## Discussion

Pursuing a new drug candidate for a disease has been exhaustively overwhelming. Therefore, drug repurposing has recently gained significant interest. Here, we present our study of enriching existing KGs with drug synergy data to achieve a primary goal: repurposing predicted drug synergy partners as single agents for the disease of interest. The alignment between the model’s predictions and the real-world literature highlights the model’s effectiveness in identifying clinically relevant and potentially impactful drug candidates. Although synergy predictions followed by safe drug selection, and pathway in pharmacome tracing, have reduced the number of selected drug candidates, they form a strong foundation for further research on these candidates.

Some of our selected candidates such as albendazole and miconazole, mefloquine, ciprofloxacin, and moxifloxacin for AD, have a robust profile of clear pathways in our pharmacome with the disease of interest as well as experimental (in cell and animal models) and clinical studies that support their willingness to be repurposed for that disease. Another portion of selected drugs, including disulfiram, auranofin, finafloxacin, taribavirin, and ivermectin for AD, has strong literature evidence but unclear pathways in our pharmacome, which require further analysis and understanding of the relation encompassed in their pathway in the pharmacome. Finally, drugs that have no literature evidence or clear pathways were marked as the least suitable for repurposing including atovaquone for AD and pyrimethamine for Schizophrenia. In addition to these groups exists a controversial group in which literature supports their harmful effect on the selected disease; however, they appear many times in our predictions as a valuable agent for controlling that disease. This group includes mefloquine, chloroquine, and albendazole for schizophrenia. We highly recommend further clinical and experimental investigation of these drugs for that disease before the commencement of their repurposing procedures.

To challenge our model’s applicability, we selected a drug for COVID-19 even knowing that there is no disease node for COVID-19 in our pharmacome. The results showed its ability to predict synergy and repurposing candidates, which a strong literature profile confirmed. This vast ability underscores that the model does not rely on a single node or relation but on the overall interaction within the network. The potential of this approach to repurposing drugs for diseases that are out of the scope of the used pharmacome gives insights into comorbidity pathways that exist between These diseases. Specifically, we refer to the possible comorbidity between COVID-19 and neurodegenerative diseases (NDD). The ability of SynDRep to find repurposing candidates for COVID-19 may be attributed to these underlying comorbidity pathways. Therefore, this work paves the way for further research already being conducted for detecting such comorbidities (COMMUTE. Comorbidity Mechanisms Utilized in Healthcare 2024).

Our study first took a broad approach that relied on graph topology metrics as well as the physicochemical properties of the drugs. ML models were then used to classify combinations into synergistic or antagonistic categories. However, this approach primarily neglected the “relationship type” factor within the KG and relied solely on the data associated with the drug nodes. In contrast, KG embedding models take these relations into account when embedding all the nodes of the KG into vectors. This consideration improves the performance and predictions of KG embedding compared to ML modeling. Therefore, this approach emphasizes the beneficial effect of organizing data into KGs and the further extraction of this data using graph embedding techniques over the classical ML approaches. Moreover, analyzing biomedical data using network structures requires a thorough understanding of network topology. Therefore, we used the topological features along with the physicochemical features of the drugs for the training and prediction in the classical ML approach. However, these methods often demand high computational and space costs (Su *et al*. 2020) and result in lower performance than the graph embedding method as evidenced by the lack of literature studies for predicted top scorer partners. On the contrary, organizing the data into a graph that can describe the complex structure of data and enables the characterization of high-order geometric patterns for the networks, improves the performance of various data analysis tasks (Xu 2020). Graph embedding techniques are able to convert sparse high-dimensional graphs into continuous low-dimensional vectors that maximally preserve the graph structure properties (Cai, Zheng and Chang 2018). The generated highly informative and nonlinear embeddings can be subsequently used for different downstream analytic tasks such as node classification and link prediction. We applied these graph embedding techniques for the prediction of the link between pairs of drugs. Unlike the ML approach, these predictions were confirmed by published scientific studies. Consequently, utilizing data represented as graphs and incorporating their embeddings represents the future direction for pharmacome data mining.

In the context of drug repurposing, maintaining a clear chain of causality from drugs to disease targets is critical. Using a broad and highly connected KG such as HBP, which contains both cause-and-effect relations and less explicit relations such as association and complexity, can lead to suboptimal model training and prediction. Although selecting a cause-and-effect subgraph is optimal, the extensive relations pool in the pharmacome captures complex protein interactions that may not be strictly cause-and-effect. Additionally, the graph needs to be large enough for effective link prediction; otherwise, performance may be compromised, which was observed with the causal-only pharmacome trial, hence pathway reviews and post-prediction literature validation were essential steps to compensate for the lack of a pure cause-and-effect subgraph by focusing on promising candidates. Another important consideration in the use of HBP is the presence of so-called “super-hubs”, which are nodes with extremely high node degrees, whose presence in the KG dilutes information and hinders learning (Sardina, Costabello and Guéret 2024). The topological imbalance in KGs has negative effects on learning using KG embedding models, where low-degree nodes embed at a much lower quality relative to high-degree nodes (Bonner *et al*. 2022). Moreover, high-degree nodes are mostly predicted as answers simply due to their higher degree, not their domain relevance (Bonner *et al*. 2022; Ratajczak *et al*. 2022). Based on these considerations, we performed a pruning of the HBP to remove non-causal relations between entities and super-hub nodes, which were mainly disease nodes. The disease nodes were removed before embedding, but we retained them in another copy of the causal-only pharmacome used for pathway confirmations. The results showed more promising repurposing candidates for AD. However, some candidates lacked pathways in the causal-only pharmacome. Additionally, we couldn’t find any connections between schizophrenia or bipolar disorder and any drugs in the pharmacome, including those usually prescribed for these diseases. Their connecting relations might have been removed during the causal relation selection step. This indicates the incompleteness of the causal-only pharmacome, which significantly impacts the repurposing approach we undertook in this study. Consequently, we recommend more manual curation of certain relation types in the HBP to enhance and update their causality comprehension. This could help maintain the relation pool present in the HBP, which is crucial for effective embedding and repurposing.

In contrast to our approach’s advantages, it exhibits some shortcomings. First, it cannot model all drug relations due to the imbalance in relation type. Most databases and studies focus on synergistic or antagonistic combinations, while scarce data about additive effect combinations are available. For instance, our work had only three additive combinations compared to tens of thousands of antagonistic or synergistic ones. Therefore, after many trials, we decided to omit these additive relations to avoid the class imbalance problem. Moreover, the embedding model’s prediction for drug combinations is not, in all cases, a drug-drug relation but may predict any other relation available in the pharmacome. Consequently, an amount of the input data might get neither synergistic nor antagonistic prediction, resulting in the loss of some combinations. Third, the controversy between some predictions and the published data about these drugs and diseases necessitates thorough investigations. Lastly, the diversity of drugs in the combination databases is limited; most are cancer-related and measure only cytotoxicity. Therefore, our approach must be extended to more specialized and highly curated KGs, such as cause-and-effect subgraphs, and additional synergistic training data.

This methodology would hold monumental potential as a robust tool for the pharmaceutical sector by broadening our search landscape and the production of more guided synergistic predictions. Moreover, this study highlights the hugely beneficial effect of computational methods not only in reducing the chemical, energy, and resource waste required to conduct thousands of wet-lab investigations but also by helping in sustainability through re-using the same drugs for more diseases and the reduction of the capital required to set up new production plans. Therefore, tons of hours, labor, and costs have been spared, which can foster further projects and speed up the pace by which treatment plans can be exploited.

## Authors’ Contributions

KSS and VSB conceived and designed the study. KSS analyzed the datasets, interpreted the results, and wrote the manuscript. BS collected data and proofread the manuscript. ATK and MHA acquired the funding. MHA, SGR, and ATK reviewed the manuscript. All authors have read and approved the final manuscript.

## Supporting information

Suplementary results and figures

## Acknowledgment

KSS gratefully acknowledge Andrea Zaliani for his insights into the manuscript.

