## Supplementary material for "SynDRep: A Knowledge Graph-Enhanced Tool based on Synergistic Partner Prediction for Drug Repurposing": Suplementary results and figures

### Results

#### 1. Classical Machine learning

A nested cross-validation was used to assess the performance of the four ML models.  As depicted in Figure S3, there is a negligible difference between the ROC-AUC scores of the four ML models. However, the elastic net model exhibited the highest ROC-AUC (0.8027) and was selected for the further prediction of synergism.

Synergy predictions between pairs of drugs in the pharmacome were carried out using the trained elastic net model. The predictions were divided based on the predicted relation between drugs into two distinct subsets: a synergy set, and an antagonism set. This categorization yielded a synergism set comprising 3,330,649 combinations and an antagonism set comprising 7,764,165 combinations. Our subsequent focus centered on the synergy set to validate the model’s predictability further. The five highest-scoring synergy combinations, in terms of synergism class probability, comprise aprinocarsen with carboxymycobactin T, salmon calcitonin, vintafolide, teniposide, or bevasiranib. They were subjected to a thorough literature review to virtually validate the reliability of the model's predictions. Although most of These drugs are used for similar indications as anti-cancers, we could not find any supporting studies validating their synergy when used in combination.

#### 2. KG embedding

The produced hyperparameters from the hyperparameter optimization (HPO) (Supplementary Table S1) were used to train the optimum model, which was then evaluated by the percentage of true predictions for all relations. RotatE was the best model to produce true predictions at the lowest rank (73.09%) as shown in Fig S4. We characterized this model results further by calculating the different models’ multi-class ROC-AUC. RotatE also produced the highest multiclass ROC-AUC (0.76), which indicated the model’s high performance in the prediction of the proper relation compared to other models (Figure S5). An additional test set exclusively containing drug-drug relations from the original test set was employed to validate RotatE as the optimal model for link prediction among drug entities within the pharmacome. The results demonstrated that RotatE consistently outperformed other models, exhibiting the highest percentage of accurate predictions at the lowest rank within this dedicated subset (61.84%), as illustrated in Figure 2.

#### 3. Synergy prediction

The set of predictions was divided based on the predicted relation between the head and tail entities, creating two distinct subsets: a synergy set (52,127 combinations) and an antagonism set (35,684 combinations). Our subsequent focus centered on the synergy set to assess the model’s predictability further and select potential drug repurposing candidates. The five highest-scoring synergy combinations are shown in supplementary Table S2.

#### 4. Drug repurposing

Based on our predictions, we selected a list of drugs that exhibited the highest score as synergistic partners with selected commonly prescribed drugs for the disease of interest. Then we chose the safe drugs and explored their pathways to the disease in the pharmacome.

##### a. Alzheimer’s candidates

To find repurposing candidates for AD, we detected the safe predicted synergistic partners with donepezil, rivastigmine, and galantamine. Mefloquine is an orally administered blood antimalarial (Palmer, Holliday and Brogden 1993) and was reported in our predicted synergy with both donepezil and rivastigmine. Moreover, Studies have shown that mefloquine significantly enhances the procognitive effect of donepezil in C57BL/6 male mice and Sprague–Dawley (8-week-old) male rats, and potentiates the donepezil-induced hemodynamic effects in the hippocampus of C57BL/6 male mice (Droguerre *et al.* 2020; Vidal *et al.* 2020). Our common pathway exploration from pharmacome revealed a pathway within our pharmacome between mefloquine and both donepezil and rivastigmine (Figure S6). Other AD repurposing candidates are ciprofloxacin, moxifloxacin, chloroquine, taribavirin, and ivermectin. They appeared in synergistic combinations with one or more drugs prescribed for AD. The common pathways in pharmacome were elucidated and some of them are shown in Supplementary Figure S7 and Figure S8, and many studies, in cell and animal models as well as clinical trials,  support their potential use for the management of AD (Zbarsky, Thomas and Greenfield 2004; Osorio *et al.* 2019; Zusso *et al.* 2019; Lindblom *et al.* 2021; Niklasson, Klitz and Lindquist 2022; Varma *et al.* 2023). Atovaquone has shown unclear pathways in our KG and no supporting published studies for its use in AD, therefore we suggest it is the least suitable for repurposing.

##### a. Schizophrenia and bipolar disorder candidates

Mefloquine stemmed from our synergy prediction with the three selected drugs for the management of Schizophrenia: olanzapine, ziprasidone, and thiothixene, although the majority of published animal models and retrospective studies demonstrate that its prolonged use induced psychosis and might worsen Schizophrenia (Alisky, Chertkova and Iczkowski 2006; Mawson 2013). The shared pathways to schizophrenia between mefloquine and the three drugs are very similar (Figure S9). Like mefloquine, the majority of published studies demonstrate that extended use of chloroquine induced psychosis and might worsen Schizophrenia (Alisky, Chertkova and Iczkowski 2006; Biswas, Sen and Majumdar 2014). However, it stems from our synergy prediction with ziprasidone and thiothixene (Supplementary Figure S10). On the other hand, pyrimethamine, an antiparasitic drug typically used to treat malaria and toxoplasmosis (Wishart *et al.* 2018), has no existing studies investigating the pure potential effect of pyrimethamine on Schizophrenia. However, most of them are discussing it in the context of reducing the symptoms of toxoplasmosis-induced schizophrenia (Webster *et al.* 2006; Castaño *et al.* 2022). The presence of pyrimethamine as a synergistic drug with ziprasidone and thiothixene highlights the need for further research on the underlying mechanisms and pathways for its use in the control of schizophrenia, even in the absence of Toxoplasmosis.

#### 5. Causal-only pharmacome

miconazole and albendazole have emerged as promising candidates for repurposing to treat AD. A prominent pathway of miconazole in association with donepezil and rivastigmine, and galantamine to AD involves the role of these three drugs as inhibitors of acetylcholinesterase (ACHE). ACHE leads to an increase in amyloid-beta precursor protein (APP) through increasing the abundance of beta-amyloid. This elevation in APP has been linked to an increased AD. Miconazole, on the other hand, acts as an inhibitor to nitric oxide synthase 2 (NOS2). NOS2 also leads to an increase in APP through increasing the abundance of nitric oxide. Since all drugs are inhibitors, they inhibit these pathways resulting in protection from AD or slowing its progression.

Albendazole shares a shared pathway to AD with donepezil and rivastigmine. As previously mentioned, the donepezil and rivastigmine pathway involves the indirect inhibition of the abundance of beta-amyloid.  In addition to increasing APP, beta-amyloid typically increases phosphorylated MAPT which is a hallmark of AD progression. Therefore, the reduction in beta-amyloid leads to inhibition of phosphorylated MAPT. On the other hand, Albendazole inhibits Tubulin alpha-1A protein (TUBA1A), leading to a decrease in Cell division control protein 42 homolog (CDC42). The decrease in CDC42 will result in a decrease in a composite of mitogen-activated protein kinase 1 and 3 (MAPK1 and MAPK3), which will eventually decrease the phosphorylated MAPT.

### Tables

**Table S1. The hyperparameter levels produced from the hyperparameter optimization process and used during the training of the optimum embedding models.**

| **Parameter** | **TransE** | **TransR** | **RotatE** | **ComplEx** | **HolE** |
| --- | --- | --- | --- | --- | --- |
| Embedding dimensions | 256 | 32 | 256 | 128 | 256 |
| Relation space dimensions | - | 224 | - | - | - |
| Scoring factor norm | L_1_ | L_1_ | - | - | - |
| Optimizer | Stochastic gradient descent | Stochastic gradient descent | Adam | AdaGrad | AdaGrad |
| Learning rate | 0.018 | 0.57 | 0.005 | 0.31 | 1 |
| Training batch size | 512 | 2048 | 256 | 1024 | 256 |
| Loss | Margin ranking loss | Margin ranking loss | Self-adversarial negative sampling loss | Softplus loss | Margin ranking loss |
| Self-adversarial sampling temperature | - | - | 0.7 | - | - |
| The margin between positive and negative scores | 3.5 | 0.5 | - | - | 1.5 |

**Table S2. The five top scorer predictions, based on the original HBP, and their validation from published studies.**

| **Rank** | **Drug A** | **Drug B** | **References** | **Remarks** |
| --- | --- | --- | --- | --- |
| 1 | Everolimus | Doxorubicin | (O’Reilly *et al.* 2011; Kim *et al.* 2016) | Studies showed an additive or a synergistic effect between the two drugs |
| 2 | Cytarabine | Fluorouracil | (Yu *et al.* 2022) | Cocrystal of cytarabine with 5-fluorouracil having synergistic antitumor effects |
| 3 | Mebendazole | Auranofin | (Partridge *et al.* 2017) | There is no study for synergy |
| 4 | Disulfiram | Cytarabine | (Bista *et al.* 2017; Yang *et al.* 2020) | studies show the two drugs are routinely used in combination but no specific study on synergism |
| 5 | Doxorubicin | Cytarabine | (Fountzilas, Inoue and Ohnuma 1990) | combination showed additive effect |

**Table S3. Validation of top scorer predictions, based on causal-only pharmacome,  from published studies.**

| **Drug A** | **Drug B** | **References** | **Remarks** |
| --- | --- | --- | --- |
| Tamsulosin | Ruxolitinib | (Wishart *et al.* 2018) | Tamsulosin decreases ruxolitinib excretion rate, which could result in a higher serum level. |
| Ruxolitinib | Zolpidem | (Wishart *et al.* 2018) | Zolpidem decreases the metabolism of Ruxolitinib increasing its effect. |
| Ruxolitinib | Prednisolone | (Cortés *et al.* 2019) | The combination of Ruxolitinib with Prednisolone showed synergistic effects. |
| Ruxolitinib | Cisapride | - | No study was found on their combination. |
| Deslanoside | Ruxolitinib | - | No study was found on their combination. |

### Figures

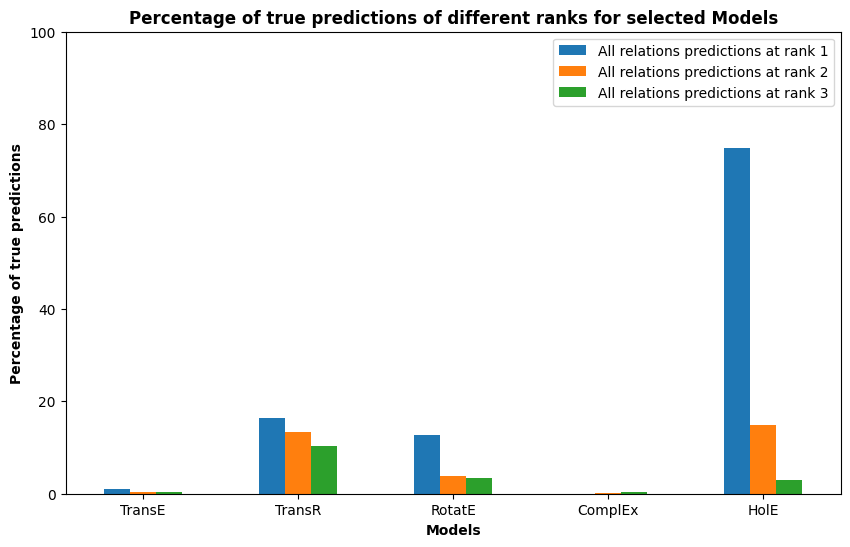

**Figure S1. Percentage of true all relations predictions at different ranks for selected models used to embed Casual-only pharmacome.** Optimum models were used to predict the test set, and then the predicted relations were compared to the actual relations to calculate the percentage of true predictions.

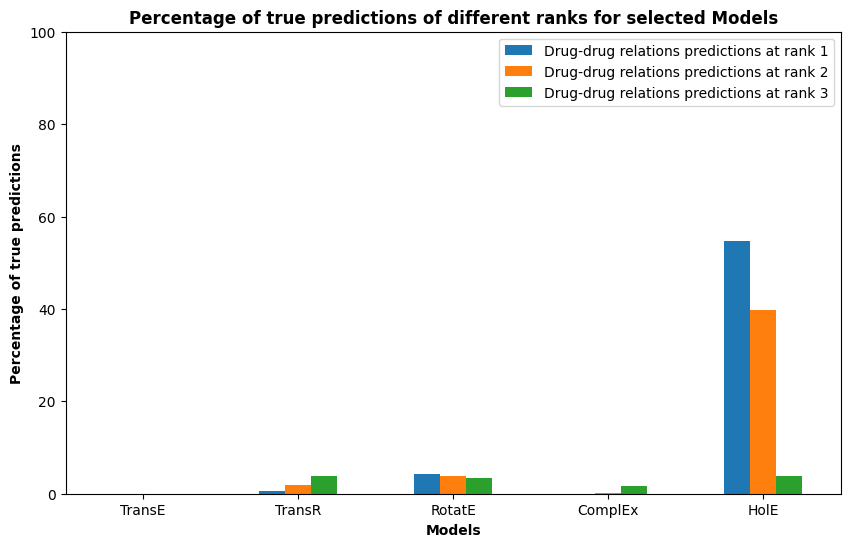

**Figure S2. Percentage of true drug-drug relation predictions at different ranks for selected models used to embed Casual-only pharmacome.** Optimum models were used to predict the drug-drug relations in the test set. Then, the predicted relations were compared to the actual relations to calculate the percentage of true predictions.

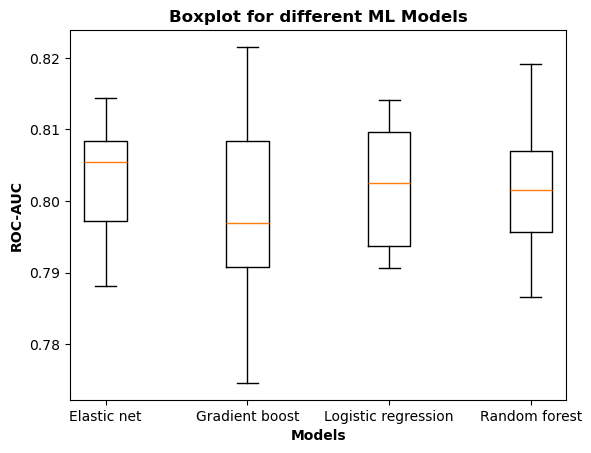

**Figure S3. Benchmarking of four ML models trained to classify between synergism and antagonism.** Each boxplot shows the distribution of the ROC-AUC values over ten repeats of the ten-fold nested cross-validation procedure.

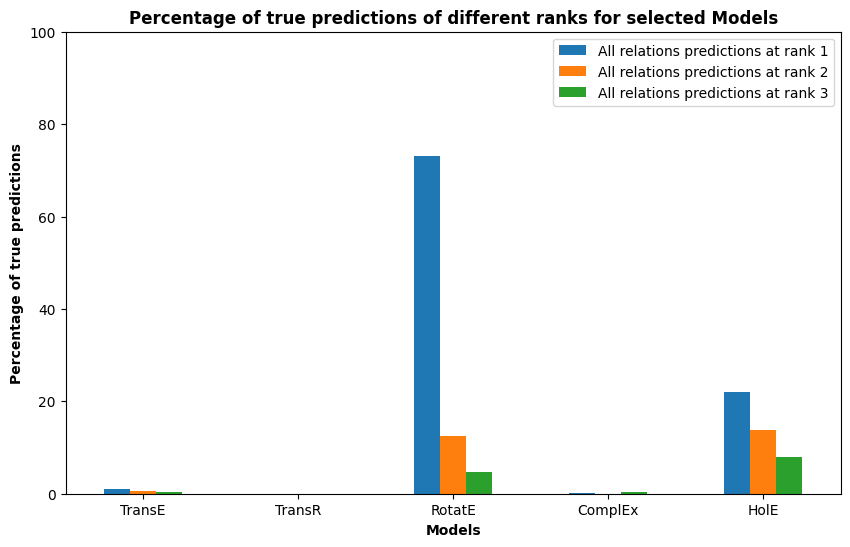

**Figure S4. Percentage of true predictions for all relations at different ranks for selected models.** Optimum models were used to predict the test set from the original HBP, and then the predicted relations were compared to the actual relations to calculate the percentage of true predictions.

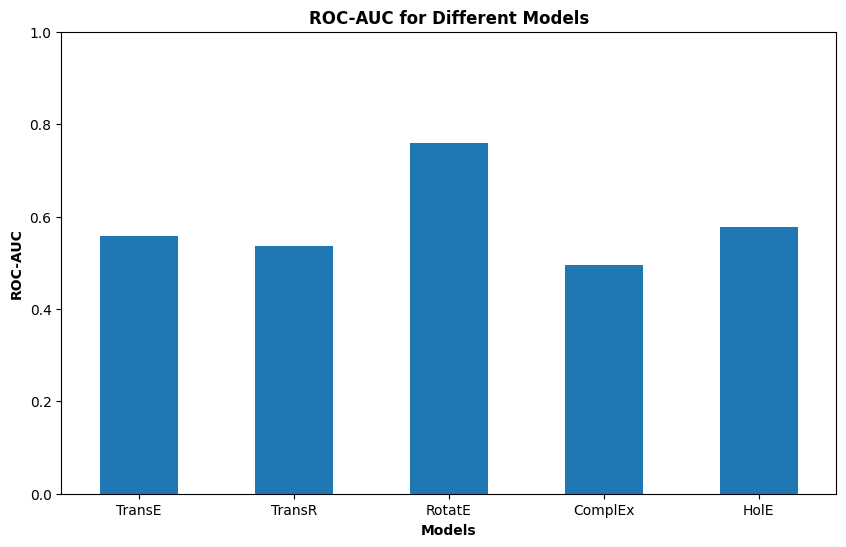

**Figure S5. Multi-class ROC-AUC for all relations predictions at rank 1 for selected model.**

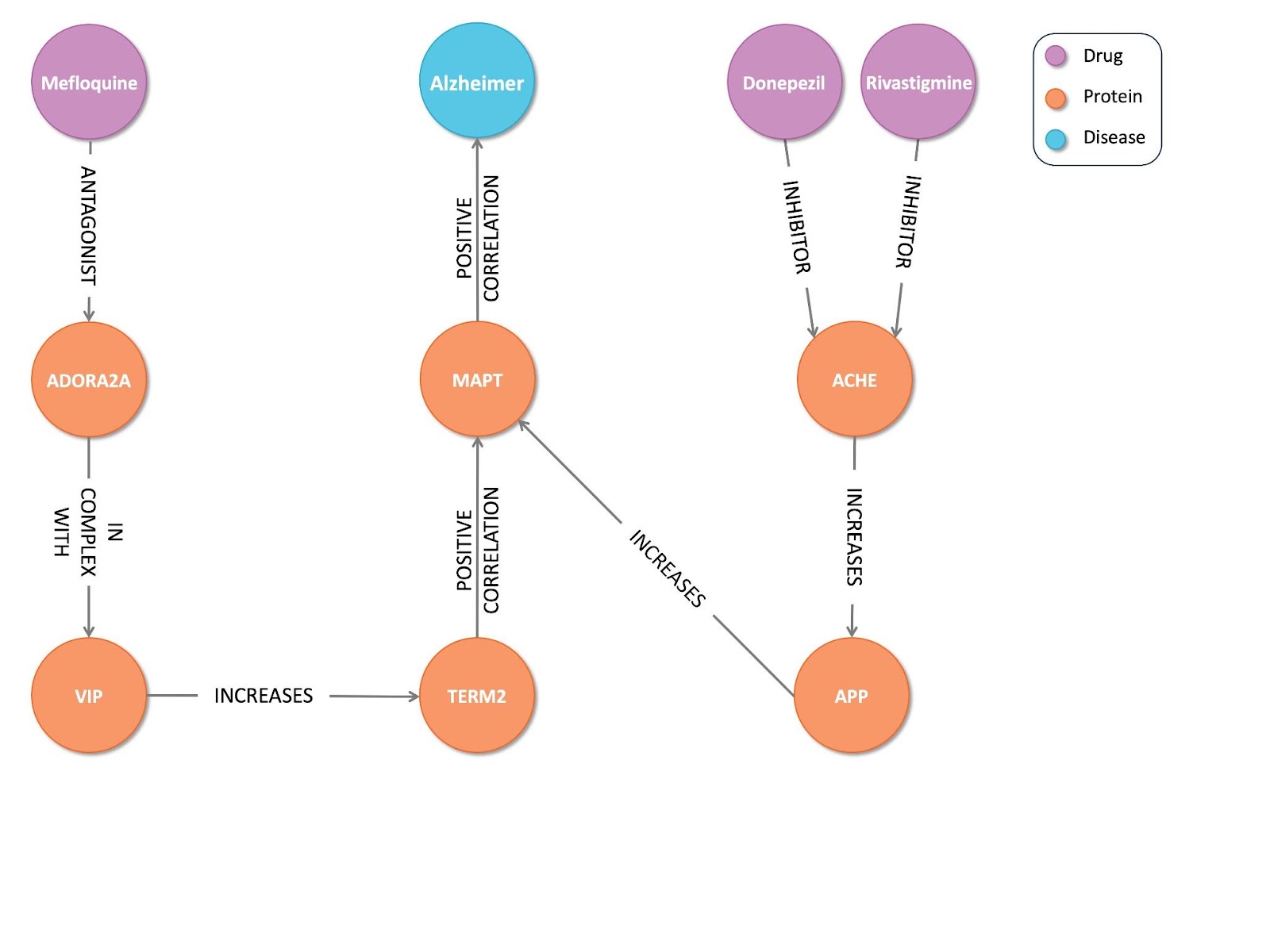

**Figure S6. The shared pathway between mefloquine and both donepezil and rivastigmine to Alzheimer’s disease.**

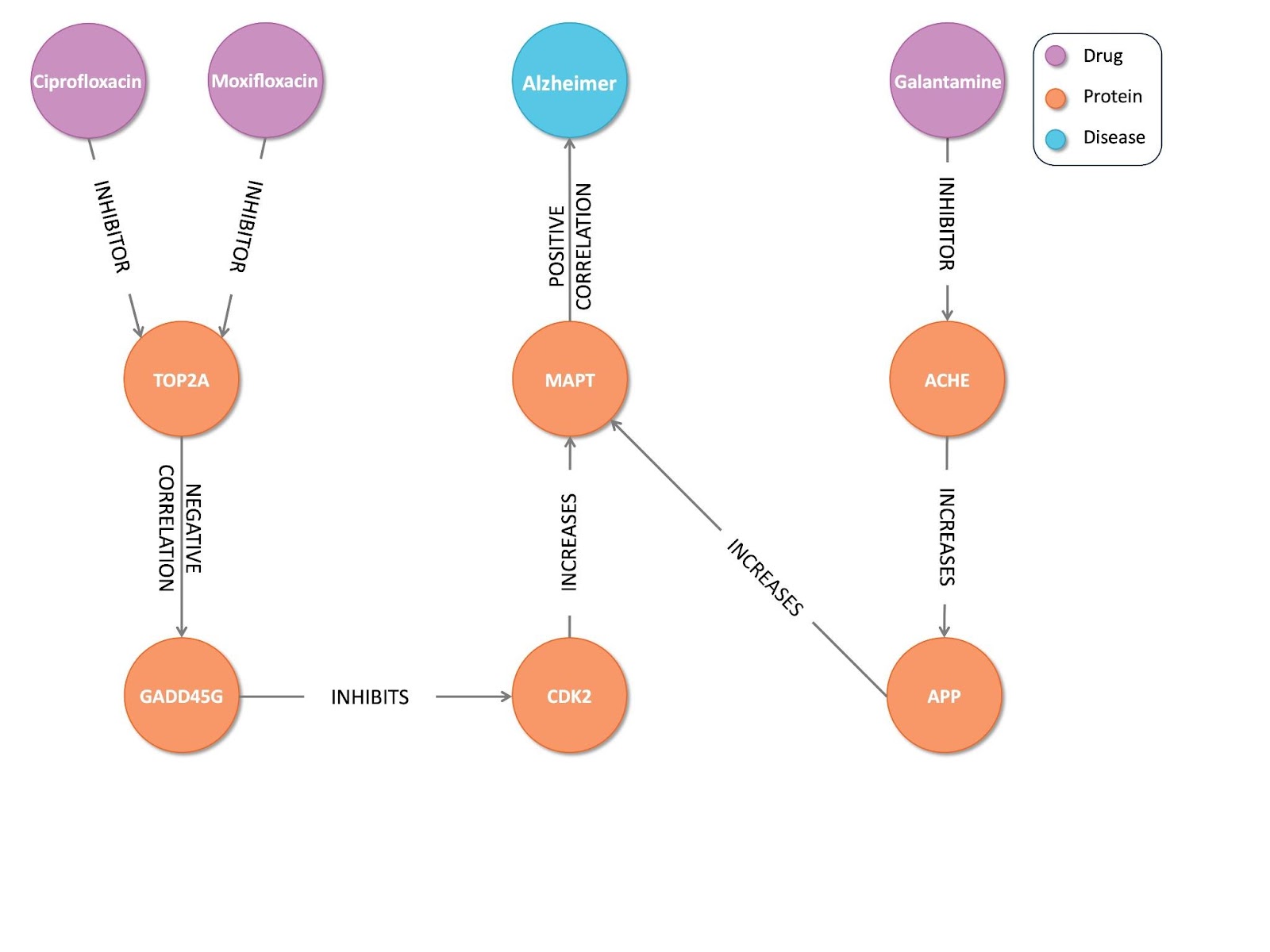

**Figure S7. The shared pathway between galantamine and both ciprofloxacin and moxifloxacin to Alzheimer’s disease.**

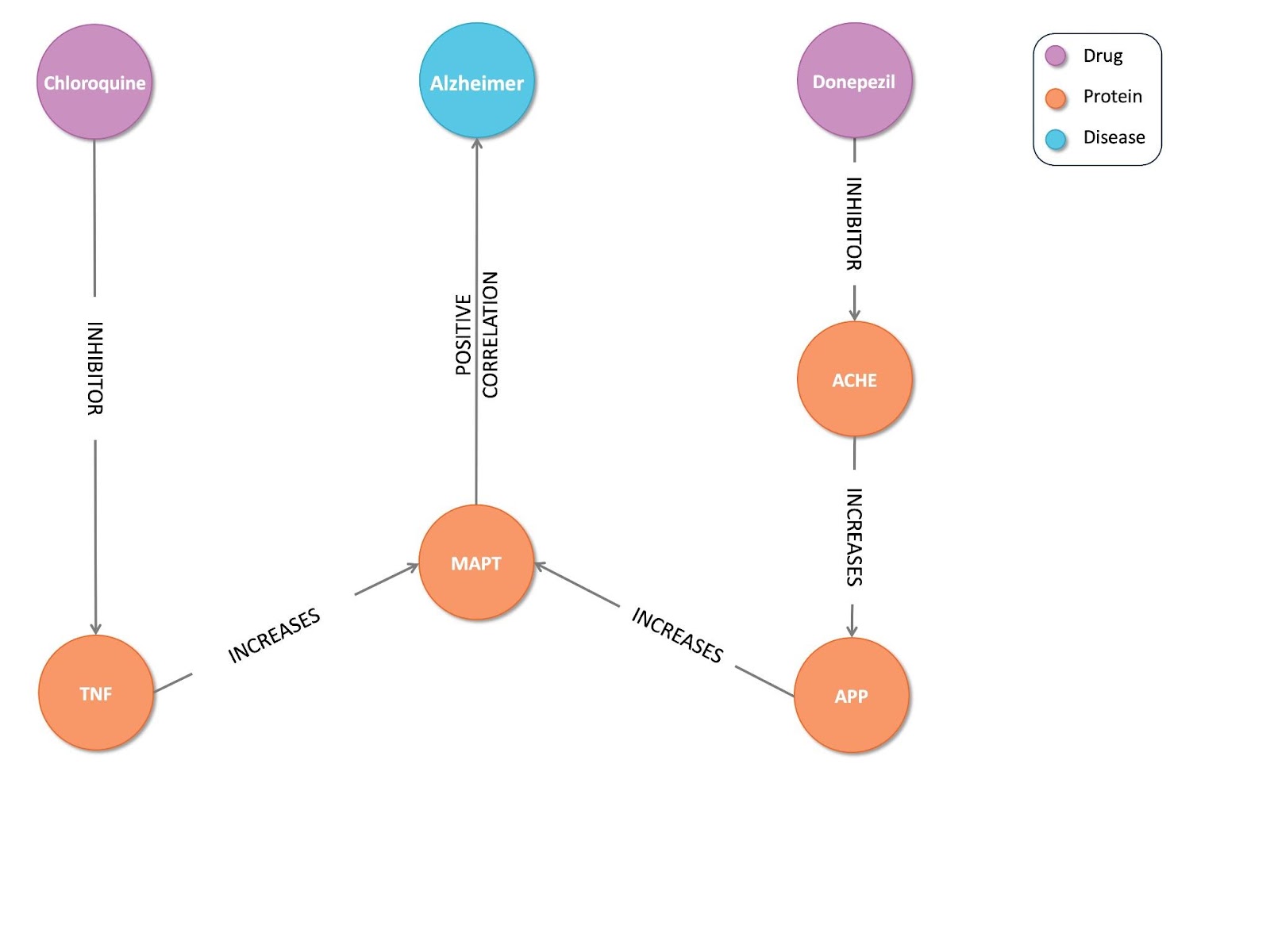

**Figure S8. The shared pathway between donepezil and chloroquine to Alzheimer’s disease.**

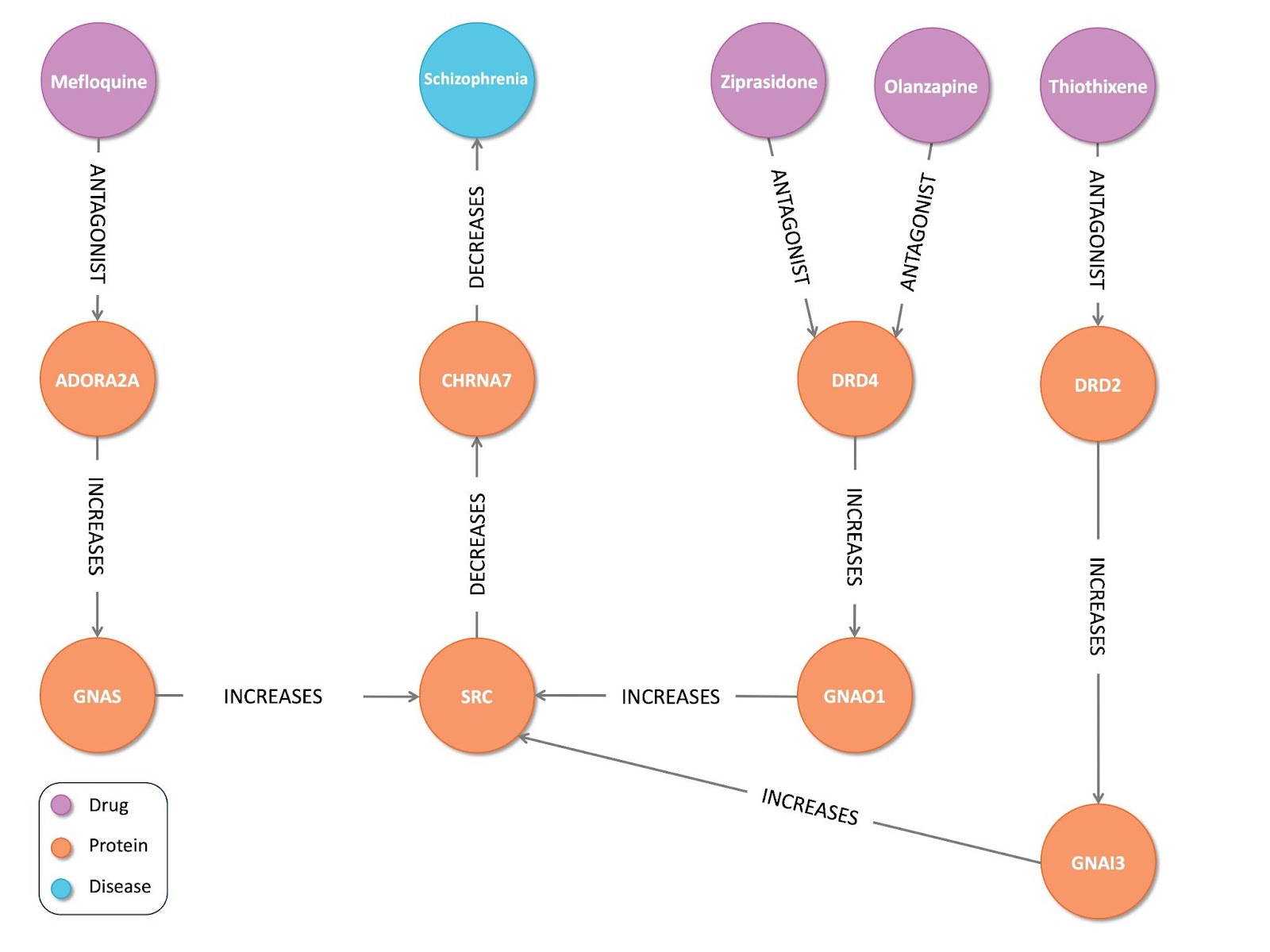

**Figure S9. The shared pathways between mefloquine, olanzapine, ziprasidone, and Thiothixene to Schizophrenia.**

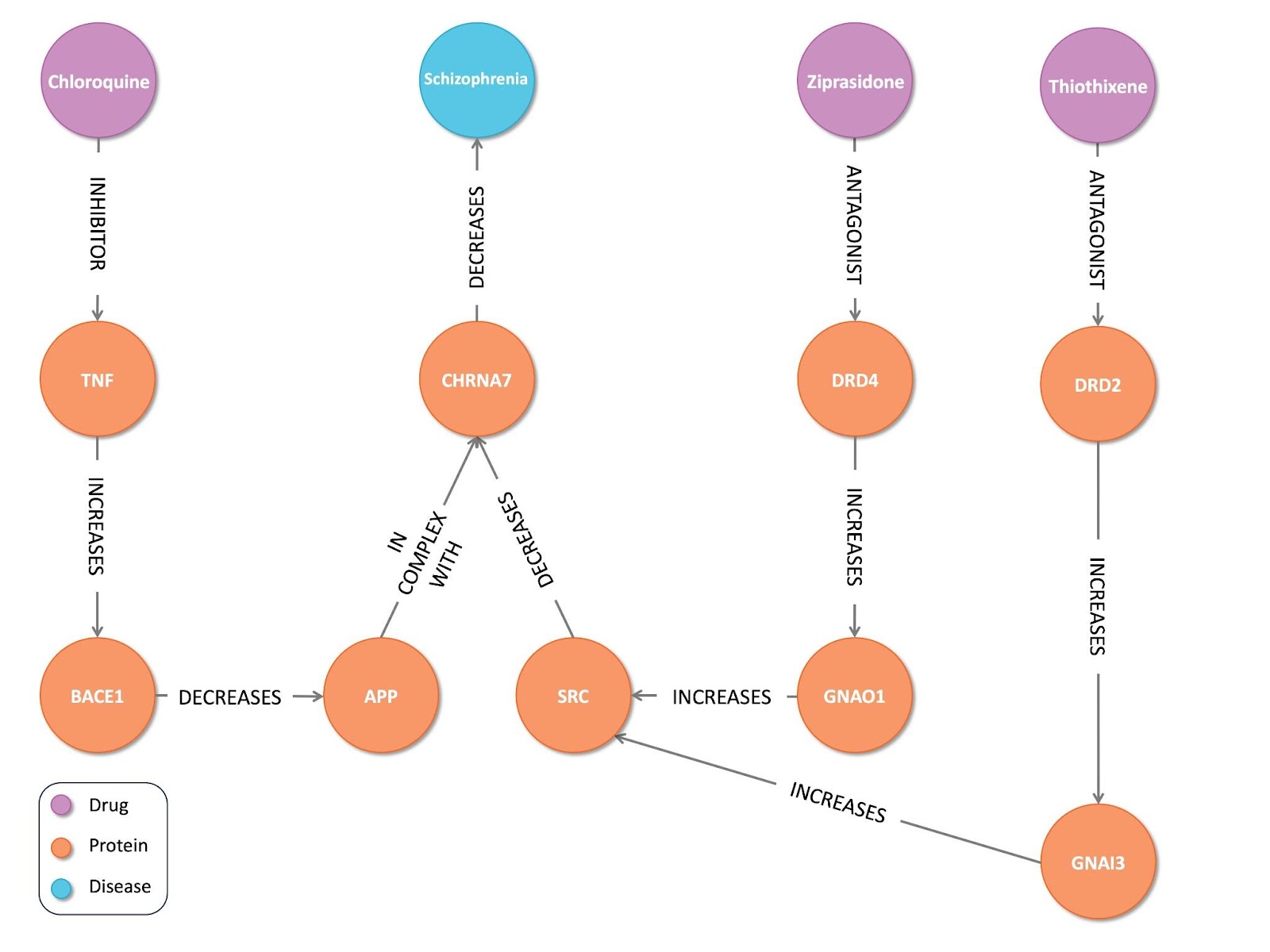

**Figure S10. The shared pathways between chloroquine, ziprasidone, and Thiothixene to Schizophrenia.**
